# The genome of *Doleromyrma darwiniana* and annotations for five additional Dolichoderinae species

**DOI:** 10.1101/2025.11.13.688362

**Authors:** Phoebe Keddell, Joseph Guhlin, Peter K. Dearden

## Abstract

Darwin’s ant, *Doleromyrma darwiniana*, is a native ant to Australia that has established invasive populations in nearby Aotearoa, New Zealand. Darwin’s ant belongs to the Dolichoderinae subfamily of the Formicidae family, which contains species with a broad range of invasion histories, including the internationally prolific Argentine ant (*Linepithema humile*). The genome of Darwin’s ant, *Doleromyrma darwiniana*, is presented here, created from a combined sample of 10 female workers. Generated using Oxford Nanopore Technologies PromethION 2 sequencing data, the assembly has a total length of 238.80 Mb and an N50 of 14.23 Mb. The assembly is compared to that of *L. humile* and four additional species from the same subfamily, the Dolichoderinae. Annotations were generated for all six Dolichoderinae species and sorted into orthogroups to assess the diversity and conservation of the Dolichoderinae genomic repertoire.

## Background and Summary

The species *Doleromyrma darwiniana* (Forel 1907)^1^, commonly referred to as Darwin’s ant, is native to Australia, particularly its coastal regions, and has successfully established invasive populations across Aotearoa, New Zealand via multiple introductions at several ports, first recorded in New Zealand in 1980^2–5^. Young *D. darwiniana* queens typically found colonies in dry, forested areas within the soil, with a preference for sites under logs or large stones. They typically reach group sizes of several hundred monomorphic workers, each approximately 2 millimetres in length^3,6^. *D. darwiniana* also frequently establishes nests in or near urban areas, earning its other common name, the brown house ant, and is known to produce a distinctive odour upon worker death or colony establishment in a household^2,6^.

The taxonomic position of *D. darwiniana* within the tribe Leptomyrmecini suggests a close relation between *D. darwiniana* and the internationally invasive Argentine ant (*Linepithema humile*). While *L. humile* has progressively extended its range to colonise over 50 countries across six continents, *D. darwiniana* has a more specialised invasion history consisting of the repeated colonisation of one country despite numerous other international possibilities facilitated by the Australian maritime industry^7,8^. Recently, four additional Dolichoderinae genomes were made publicly available on the National Centre for Biotechnology Information (NCBI) database by the Global Ant Genomic Alliance. Two of these species (*Iridomyrmex anceps* and *Ochetellus glaber*) belong to the same tribe as that containing *D. darwiniana* and *L. humile*, the Leptomyrmecini. In contrast, the remaining two species (*Liometopum microcephalum* and *Tapinoma melanocephalum*) belong to a different tribe, the Tapinomini. These species span a wide range of invasive efficiency (Table 1), providing a strong comparative dataset. This study helps to build the foundation for such a comparison by generating new protein annotations for all six Dolichoderinae species to assess the extent of conservation and genomic diversity across the subfamily. The addition of *D. darwiniana* signifies the least invasive and/or internationally widespread member of the Leptomyrmecini tribe sequenced to date (Table 1).

**Table 1:** The native and invasive range of each Dolichoderinae species is included. Species are considered to be invasive ‘worldwide’ if they have successfully colonised all or all but one continent they are non-native to; this does not imply they have colonised every country. Species’ invasiveness is considered low, intermediate, or high: Low invasiveness species have colonised a maximum of one non-native country, highly invasive species are those established worldwide, and intermediate species fall between these two categories. Species ranges are identified using AntMaps (https://antmaps.org/).

| Species | Tribe | Native Range | Invasive range | Invasiveness |
| --- | --- | --- | --- | --- |
| *Doleromyrma darwiniana* | Leptomyrmecini | Australia | Aotearoa, New Zealand | Low |
| *Iridomyrmex anceps* | Leptomyrmecini | Australia to Southeast Asia | Iran, Yemen, Society Islands | Intermediate |
| *Linepithema humile* | Leptomyrmecini | Central South America | World-wide | High |
| *Ochetellus glaber* | Leptomyrmecini | Australia to Southeast Asia | Several Pacific Islands, Florida, South Africa, New Zealand | Intermediate |
| *Liometopum microcephalum* | Tapinomini | Central Europe to Iran | None confirmed | Low |
| *Tapinoma melanocephalum* | Tapinomini | Southeast Asia | World-wide | High |

## Methods

### *Doleromyrma darwiniana* Sample Details

Samples of workers from the same nest were collected by Richard Toft in Nelson, New Zealand (Latitude: −41.16530000, Longitude 173.16360000) on the 17^th^ of August 2024 and stored at −20°C in 70% ethanol before DNA extraction. DNA extractions were performed on individuals and pools of two, five, and ten diploid females, with the largest pool of 10 being selected to be used for library preparation due to the small amount extracted from pools of fewer individuals. Given the small size of the specimen, pooling to gain enough DNA was required. Although *D. darwiniana* mating biology has not received specific study, pooling was thought to be acceptable given that the species within the Dolichoderinae tend to be monogynous and monandrous, indicating the pooled sample consisted of full sisters^9,10^. The effect of pooling is further minimised as the source colony is presumed to have experienced a population bottleneck due to its recent invasion history^11^.

### *Doleromyrma darwiniana* Genomic DNA Extraction

Genomic DNA was extracted from the pooled sample in line with the *Quick*-DNA Magbead Plus Kit protocol for solid tissue extractions by Zymo Research, with some minor adjustments to maximise DNA yield (Zymo Research, D4081). Briefly, before the incubation at 55ºC of the sample with Proteinase K, the sample was agitated via a sterilised pestle to homogenise the pooled sample with the fluid, increasing Proteinase K access to the sub-exoskeletal tissues containing the majority of the ants’ genomic DNA. After agitation, the incubation was extended from the suggested one to three hours to approximately 16 hours. Additionally, throughout the post-incubation DNA purification process, the period allowed for DNA to bind to the kits accompanying MagBeads was extended from 10 to 15 minutes. Similarly, the elution period to extract the genomic DNA held on MagBeads into the associated DNA elution buffer was extended from 5 to 15 minutes and performed at 37 ºC instead of the recommended room-temperature elution. The final elution was then analysed via Agilent ScreenTape System to assess the initial size distribution of the extracted genomic DNA (Agilent, D1000) and its concentration quantified using the Qubit 1X dsDNA HS Assay Kit (Invitrogen, Q33231) to report a final extraction concentration of 17.6ng/µL in a volume of 47µL, implying a total yield of 827.2ng of genomic *D. darwiniana* DNA to be taken forward for library preparation.

### *Doleromyrma darwiniana* Library Preparation and Sequencing

The genomic library was prepared following the instructions of the Oxford Nanopore Technologies Ligation Sequencing DNA V14 and accompanying reagent kit (Oxford Nanopore Technologies, SQKLSK114; GDE_9161_v114_revW_29Jun2022). Similar minor adjustments to those made throughout the extraction process were performed to maximise the final yield of genomic DNA, namely the incubation period allowing DNA to bind to AMPure XP beads, and the elution period to return the DNA to solution throughout the DNA repair and end prep were extended from five and two minutes respectively to ten, with the elution also being conducted in a 37 ºC incubator. This temperature adjustment was additionally conducted during the final elution of adaptor ligation and clean-up, in which it was explicitly suggested. The retained library was again quantified via Qubit 1X dsDNA HS Assay Kit (Invitrogen, Q33231) to return a final concentration of 13.7ng/µL in 24µL, implying a final total of approximately 328.8ng input DNA to be sequenced. The library was sequenced on a PromethION2 flow cell (Oxford Nanopore Technologies, R10.4.1, FLO-PRO114M).

### *Doleromyrma darwiniana* Genome Assembly and Processing Methods

Once sequenced, reads were duplex basecalled using Dorado v0.8.2, filtered based on minimum phred score (10) and length (500bp) using Chopper v0.9.0^12^, and assembled with Flye v2.9.5 using the high-quality (hq) mode^13^. The repetitive regions of the assembly were soft-masked using RepeatModeler v.2.0.6 (including the long-terminal repeat pipeline) and RepeatMasker v.4.1.8 using the NCBI engine and xsmall setting to return masked regions in lowercase^14^. Assembly statistics and completeness for *D. darwiniana* in relation to the rest of the available Dolichoderinae genomes were assessed using Assembly-Stats v.1.0.1, BBMap v.39.33 and BUSCO (Benchmarking Universal Single Copy Orthologs) v.6.0.0 using the hymenoptera_odb12 dataset^15,16^. Blobtoolkit v.4.4.6 was used to visualise characteristics of the *D. darwiniana* assembly^17^.

### Annotation Generation and Orthogroup Identification

The remainder of assemblies were downloaded from GenBank^18–22^. The assemblies of all six Dolichoderinae species were filtered to remove scaffolds under 1kb and annotated using Braker v.3.0.8^15,23–35^. This filtration process is intended to minimise the inclusion of any erroneously assembled contigs, and only removed one contig from each of two species: *D. darwiniana* itself and *Ochetellus glaber*. Annotations were generated using DIAMOND to filter out redundant genes during AUGUSTUS training, protein hints from the available protein annotations of the Hymenopteran model organism *Apis mellifera* (the Western Honeybee) downloaded from NCBI (Assembly ID: Amel_HAv3.1, RefSeq ID: GCF_003254395.2), and the hymenoptera_odb10 BUSCO dataset^32,36^. Alternative transcripts were removed to retain only the primary transcripts, and the resultant protein set was assessed for completeness using the hymenoptera_odb12 BUSCO protein dataset.

OrthoFinder v.3.0.1b was used to assign annotations across all species into orthogroups using the Multiple Sequence Alignment method of gene tree and species tree inference (-M msa). A second OrthoFinder analysis incorporating annotations including annotations processed in the same manner from a species from the most closely related subfamily of the Dolichoderinae (*Pseudomyrmex concolor* from the Pseudomyrmecinae subfamily^37^) as an outgroup for accurate rooting of the phylogeny^38–41^. Orthogroup distribution was visualised in RStudio using ggplot2 and the resultant phylogenetic tree pictured using TreeViewer v.2.2.0^42,43^.

### Data Records

Sequencing and assembly outputs of this study are stored with reference to the BioProject accession PRJNA1346102. Raw sequencing data is stored on the Sequence Read Archive (SRA) under the accession number SRR35808540. Annotations are deposited in GTF format on Zenodo, as are the files related to OrthoFinder analyses (A compressed folder of individual orthogroups in FASTA format and a CSV file of copy number counts of every species for every orthogroup) both stored under the DOI 10.5281/zenodo.17420836. The final *D. darwiniana* assembly is submitted to GenBank and will be stored on Zenodo under the same DOI submission prior to the completion of GenBank processing^44^.

### Data Overview

#### Genome Report

General assembly statistics and of the *D. darwiniana* genome are visualised in a snail plot (Figure 1) and described in Table 2, alongside the equivalent metrics for the filtered genomes of the other Dolichoderinae species. In all measurements, the *D. darwiniana* genome appears to be in alignment with the typical Dolichoderinae assembly (Table 2). The final 238.8Mb genome contains no gaps between contigs, and 97.48% of the assembly is contained within contigs over 50Kb. Retroelements and DNA transposons made up 4.91% and 3.07% of the total genome, respectively, encompassed within the 46.76Mb or 19.59% of the assembly identified as repetitive elements and subsequently soft-masked.

**Table 2:** Genomic assembly metrics of *D. darwiniana* in comparison to other available Dolichoderinae assemblies. All assemblies are quantified after the removal of scaffolds under 1Kb.

| Species | *Doleromyrma darwiniana* | *Iridomyrmex anceps* | *Linepithema humile* | *Ochetellus glaber* | *Liometopum microcephalum* | *Tapinoma melanocephalum* |
| --- | --- | --- | --- | --- | --- | --- |
| NCBI TaxID | 609327 | 610584 | 83485 | 255795 | 611171 | 219810 |
| Assembly Identifier | Doldar_A1 | ASM4854278v1 | Lhum_UNIL_v1.0 | ASM4854365v1 | ASM4854256v1 | ASM4854234v1 |
| GenBank Accession | TBC | GCA_048542785.1 | GCA_040581485.1 | GCA_048543655.1 | GCA_048542565.1 | GCA_048542345.1 |
| Assembly length (Mb) | 238.80 | 223.69 | 244.44 | 300.04 | 230.33 | 252.02 |
| Average assembly depth | 194 | 84.32 | 58 | 59.27 | 98.66 | 118.6 |
| Number of scaffolds | 455 | 466 | 15 | 665 | 452 | 206 |
| Scaffold N50 (MB) | 14.2 | 9.6 | 31.4 | 3.6 | 1.2 | 6.2 |
| Longest scaffold (Mb) | 21.91 | 16.80 | 47.04 | 18.33 | 6.71 | 22.58 |
| Total GC content (%) | 37.19 | 36.17 | 37.22 | 37.25 | 37.78 | 38.00 |
| Genome BUSCO % (Complete) | 99.2 | 96.7 | 95.3 | 97.6 | 97.9 | 97.9 |
| Genome BUSCO % (Duplicated) | 0.4 | 0.4 | 0.7 | 0.4 | 0.6 | 0.5 |

**Figure 1:**
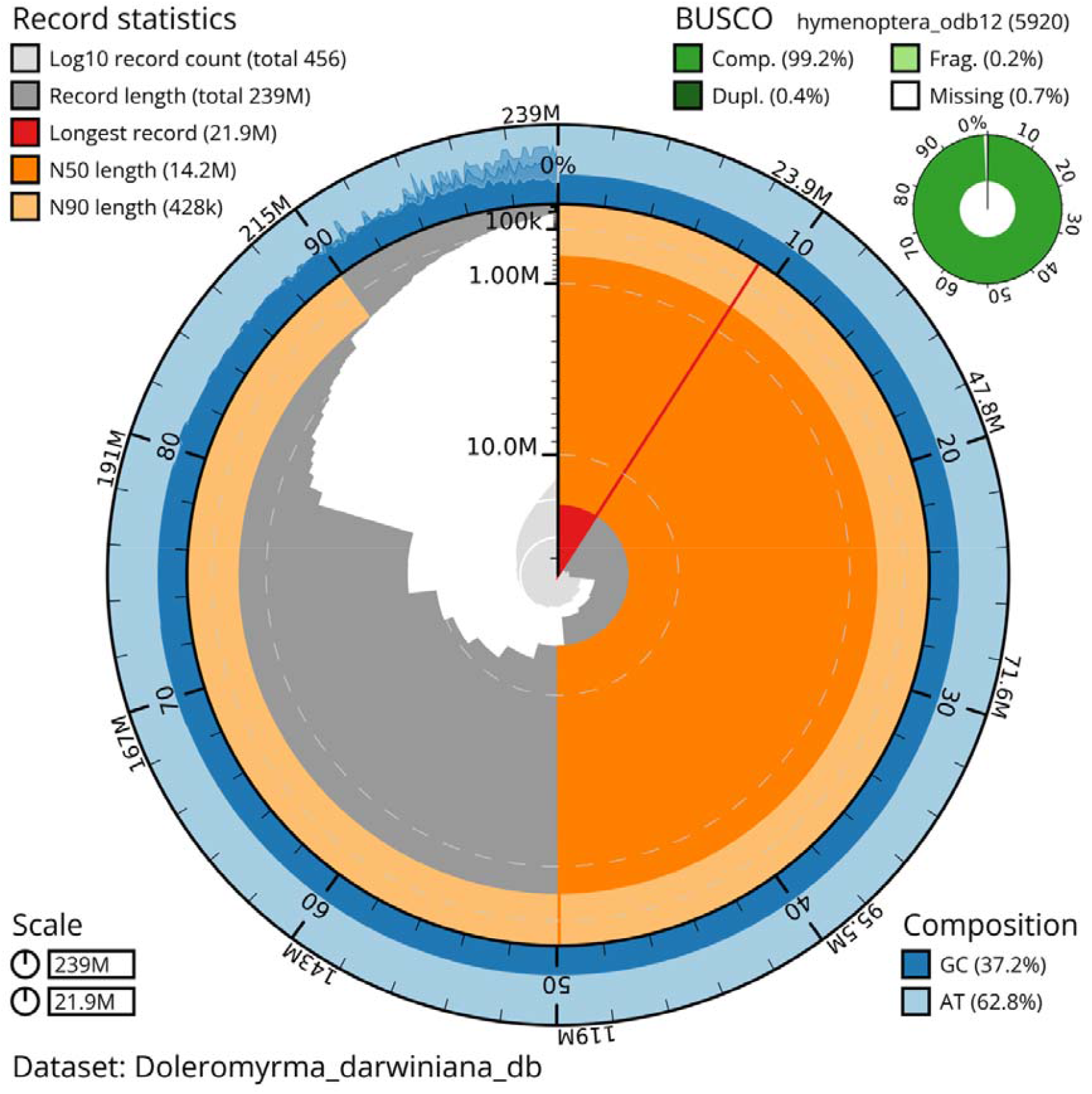
Snail plot summary of assembly statistics for the *Doleromyrma darwiniana* assembly. BlobToolKit generated snail plots that quantify several assembly metrics and BUSCO scores. The main plot is divided into 1,000 size-ordered bins around the circumference, with each bin representing 0.1% of the 238,804,658 bp assembly. The distribution of sequence lengths is shown in dark grey with the plot radius scaled to the longest sequence present in the assembly (21,915,290 bp, shown in red). Orange and pale-orange arcs show the N50 and N90 sequence lengths (14,230,604 and 427,775 bp), respectively. The pale grey spiral shows the cumulative sequence count on a log scale with white scale lines showing successive orders of magnitude. The blue and pale-blue area around the outside of the plot shows the distribution of GC, AT and N percentages in the same bins as the inner plot. A summary of complete, fragmented, duplicated and missing BUSCO genes in the hymenoptera_odb12 set is shown in the top right.

### Annotations and Orthogroups

The Dolichoderinae assemblies display a large variation in the number of identified proteins, ranging from 33,224 in *T. melanocephalum* to 69,163 in *O. glaber* (Table 3). The *D. darwiniana* genome occupies a midpoint in the Dolichoderinae range, with 43,477 identified proteins (Table 3). All Dolichoderinae annotation sets had at least a 92% complete hymenoptera_odb12 protein BUSCO score (Table 3). Despite this variation in annotation number, 85.5% of annotations could be assigned to one of 41,325 orthogroups, over a quarter of which contain an annotation belonging to each species (11,787), including 8832 entirely single-copy groups. Of the combined annotation set, 50% of annotations were in orthogroups of at least six sequences (G50: 6) and were contained within the 11,872 largest orthogroups (O50: 11872). The 11,787 orthogroups with all species present were closely followed in number by groups private to two species (Figure 2), and 4.4% of input annotations (12607 sequences) were assigned to one of 2321 species-specific orthogroups. The phylogeny recovered from the Dolichoderinae annotations (Figure 3) was in accordance with existing literature, supporting the reliability of the dataset^45,46^.

**Table 3:** Annotation Results. Counts and BUSCO evaluation of protein annotations for all six Dolichoderinae species

| Species | Annotations | BUSCO (Complete %) | BUSCO (Duplicated %) |
| --- | --- | --- | --- |
| *Doleromyrma darwiniana* | 43477 | 95.4 | 0.3 |
| *Iridomyrmex anceps* | 56375 | 92.6 | 0.5 |
| *Linepithema humile* | 41567 | 95.3 | 0.9 |
| *Ochetellus glaber* | 69163 | 93.7 | 0.5 |
| *Liometopum microcephalum* | 42570 | 93.2 | 0.6 |
| *Tapinoma melanocephalum* | 33224 | 92.1 | 0.3 |

**Figure 2:**
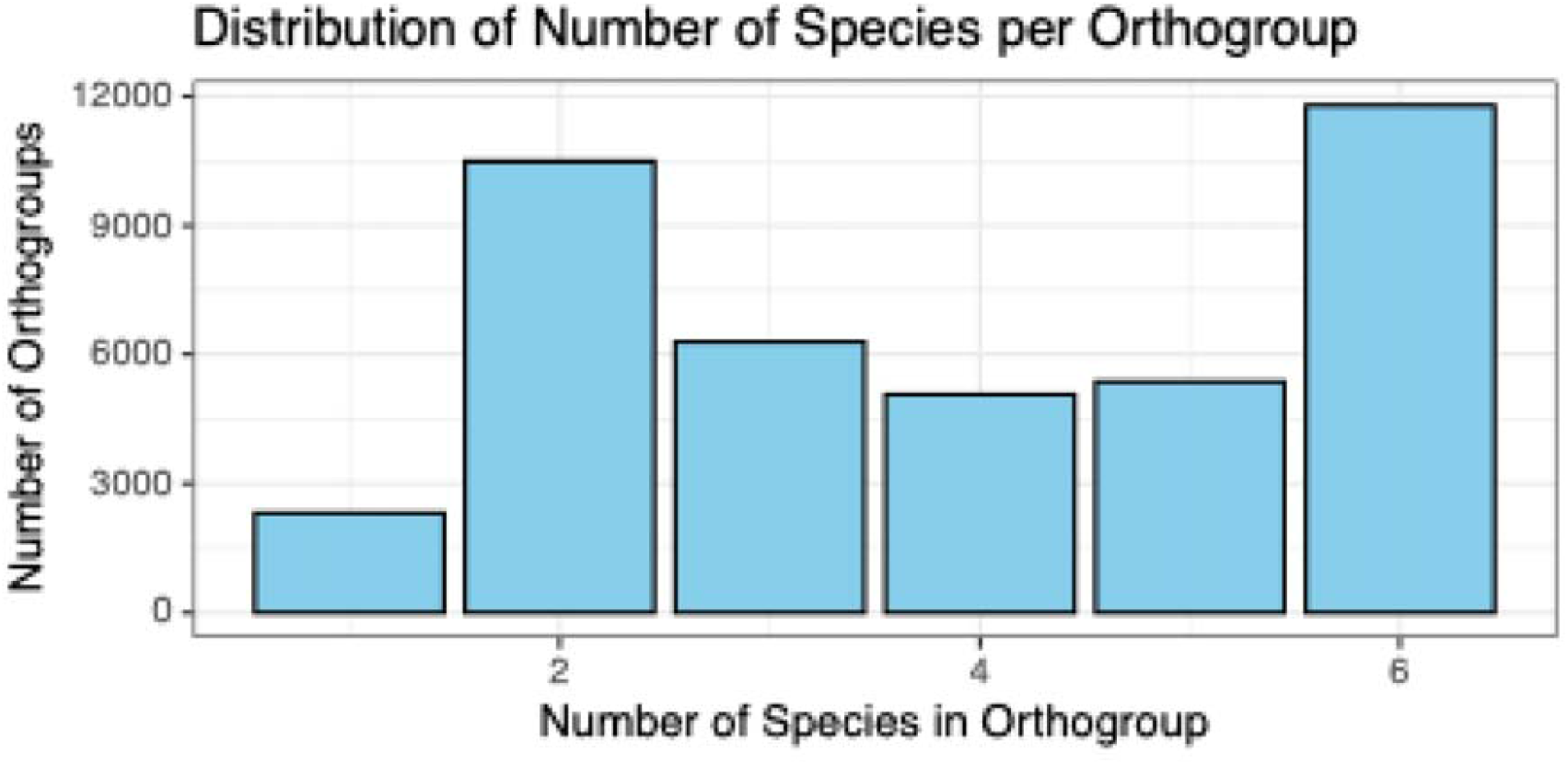
Distribution of the number of species contained in the 41,325 identified Dolichoderinae orthogroups. 1: 2321, 2: 10491, 3: 6292, 4: 5065, 5: 5369, 6: 11787.

**Figure 3:**
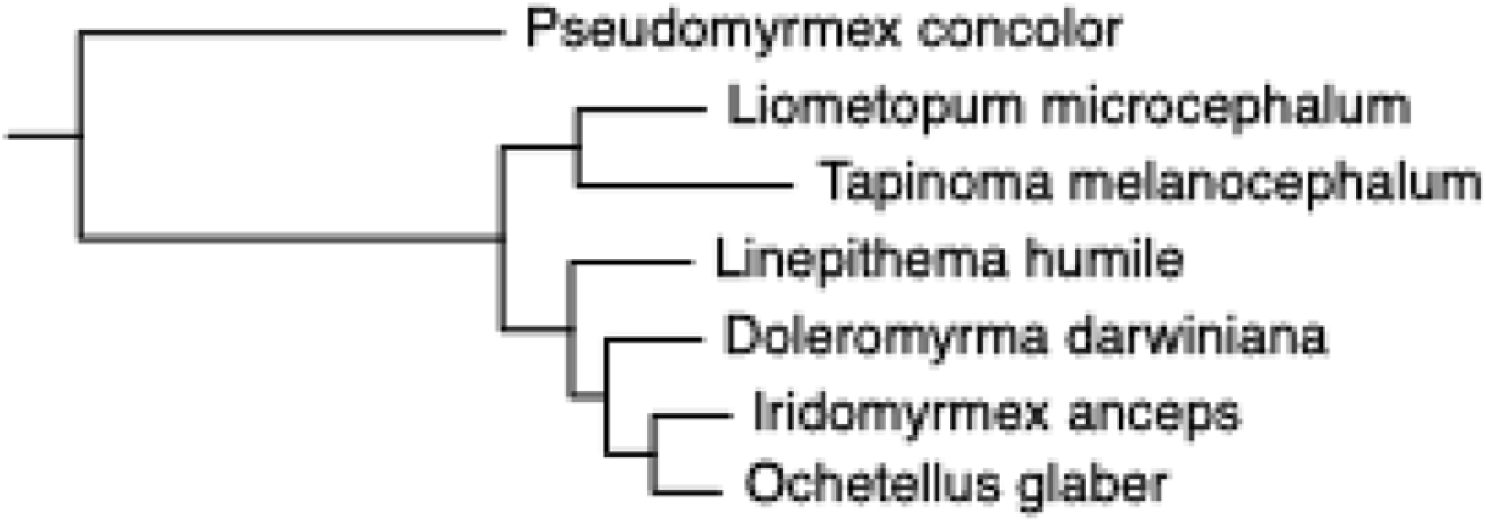
Phylogenetic tree of Dolichoderinae annotations. Tree generated as automatic output from OrthoFinder based on 6415 single-copy orthogroups identified using the Multiple Sequence Alignment (MSA) method with all seven species present.

### Summary

We present the first genome assembly for Darwin’s ant, *Doleromyrma darwiniana*, accompanied by the annotation set of all other Dolichoderinae species with available genomes on NCBI, across which a diverse range of invasion histories can be identified. Of the annotations, 85.5% were able to be assigned to an orthogroup, over a quarter of which contained all species. This provides a foundation for the further study of the Dolichoderinae subfamily at the genomic level, to address the internationally widespread effects of its members.

### Technical Validation

Before assembly, the raw sequencing data from the *D. darwiniana* extraction was filtered based on length (minimum 500bp) and phred score (minimum 10) using Chopper v.0.9.0 (Discussed in Methods) to ensure the quality of the materials used in the assembly. Several assembly metrics are compared to the available assemblies of the other Dolichoderinae species to confirm that it aligns with the results of closely related species, particularly in total length and GC content. The quality of the resultant genomic assembly for *D. darwiniana* was assessed using the hymenoptera_odb12 dataset of BUSCO (Benchmarking Universal Single Copy Orthologs) v.6.0.0. This measures the proportion of the genes expected to be present and single copy in all species of the taxon that are identifiable in the query assembly. Additionally, equivalent BUSCO analyses were performed on the assemblies of the remaining Dolichoderinae species and the annotation sets for all species. The results of the BUSCO analyses are described in the Data Overview. Further evidence towards the accuracy of the annotation sets is provided by the conservation identified throughout the OrthoFinder analysis and its recovery of an accurate phylogeny using the annotation sets.

## Data Availability

All data used within this study are submitted to NCBI under the BioProject accession PRJNA1346102. Sampling details are submitted to the NCBI in association with the BioSample accession SAMN52826755. Raw sequencing data is available from NCBI’s Sequence Read Archive (SRA; SRR35808540). The final *Doleromyrma darwiniana* genomic assembly is submitted to GenBank. Annotation sets for each species are available on Zenodo, as are the relevant OrthoFinder files and the *D. darwiniana* final assembly, which will also be available from GenBank upon the completion of processing^44^.

## Code Availability

The bioinformatic workflow, including version and parameters, is described throughout the Methods section. No independent adjustments were made to the programs discussed, and default parameters of those programs were used when not stated otherwise.

## Author Contributions

P.K.D. conceived of the study and arranged sample procurement. All authors contributed to its design. P.K. generated the *D. darwiniana* assembly, and data analysis was performed by P.K. and J.G. The manuscript draft was written by P.K. The final manuscript was revised by all authors and approved for publication.

## Acknowledgements

We wish to acknowledge Richard Toft for his contribution to sample identification and procurement, in addition to the support provided to this study by the University of Otago and Genomics Aotearoa.

